# Protective Effects of Matrine on Acute Myocardial Hypertrophy Induced by Isoproterenol via Akt/mTOR/p70S6K/eIF4E Signaling Pathway in Rats

**DOI:** 10.1101/2021.01.15.426879

**Authors:** Hui Ma, Hongwan Dang, Shijie Wei, Xiaoying Yang, Wenping Zhang

## Abstract

This study aimed to investigate whether matrine (Ma) attenuates isoproterenol (ISO)-induced acute myocardial hypertrophy via activating Akt/mTOR/p70S6K/eIF4E signaling pathway in rats. We collected 42 male Sprague–Dawley rats weighing 300±20g, randomly divided into seven groups (n=6). The myocardial hypertrophy (MH) model was well established by 85 mg/kg ISO. Changes in hemodynamic parameters were recorded using electrocardiogram after 24h with ISO injection. Western blot and real-time polymerase chain reaction were used to evaluate the expression of Akt, mechanistic target of rapamycin (mTOR), p70S6K, and eIF4E. Tissue arrangement of the 200 and 100 mg/kg Ma+ISO groups was regularly smaller than that of the ISO group. For the protein expression, Akt values in the 200 and 100 mg/kg Ma+ISO groups were 1.57- and 1.81-fold higher than in the ISO group, respectively. Moreover, compared with the ISO group, the expression trends of mTOR in the 200, 100, and 50 mg/kg Ma+ISO groups significantly downregulated. The levels of p70S6K and eIF4E reduced in the 200, 100, and 50 mg/kg Ma+ISO groups according to the ISO group (P<0.05). MRNA expression of p70S6K and eIF4E in the ISO group were 1.90- and 6.38-fold higher compared with that in the 100 mg/kg Ma+ISO group. Ma exerted neuroprotective effects against pachyntic injury. Akt activity was accelerated, but activities of mTOR, p70S6K, and eIF4E were inhibited by Ma. Activation of the Akt/mTOR/p70S6K/eIF4E signaling pathway might be the targets for the protective effects of Ma on acute myocardial hypertrophy in rats.

## Introduction

The myocardium requires a great amount of oxygen and nutrients, and insufficient blood supply causes acute myocardial hypertrophy (MH) to the heart. MH is one of the serious heart diseases, leading to relatively high morbidity and mortality [1]. The World Health Organization showed that MH is the leading cause of death, severely influencing the quality of life [2]. The primary features of MH are decreasing blood pressure, hypoxia, insufficient blood flow, and pectoralgia [3,4]. Although essential progresses in basic and clinical studies have been achieved, fundamental breakthroughs were limited in the mechanism of MH. To date, investigating the potential mechanism of MH is greatly important.

Previous study demonstrated that although the PI3K/Akt/mTOR signaling pathway regulated mRNA expression [5], Akt phosphorylation had not altered hypertrophy [6]. Short-term Akt activation showed beneficial effects on myocardial hypertrophy via improving contractile function, whereas long-term Akt activation-induced pathological hypertrophy [7,8]. The classical cardioprotective pathway against MH injury was Akt-associated signaling pathway, and the cardioprotective effects of Akt signaling in MH are well accepted [9].

Moreover, mechanistic target of rapamycin (mTOR) is activated by Akt, which altering metabolism, and increased protein expression [10]. Cardiac mTOR phosphorylation and overexpression provided cardioprotection by enhancing recovery against MH [11]. The downstream of Akt signaling was mTOR, which was essential and necessary to protect the heart against MH injury [9]. The activation of Akt/mTOR signaling pathway in the short-term would provide cardioprotection against hypertrophy, but the long-term activation will lead to hypertrophy. Thus, the activation period is worth exploring. Activation and overexpression of Akt increased p70S6K activity, thereby exerting potent cardioprotective effects responsible for hypertrophy [12]. Inhibition of Akt and p70S6K would increase MH. Meanwhile, after acute MH, activating mTOR would provide cardioprotection by accelerating cardiac function and metabolism [13]. Discovery and development of safe and effective agents to prevent acute MH in traditional medicine are clinically significant.

Matrine (Ma) (Fig.1) as a classical quinolizidine alkaloid is extracted from the roots of *Sophora flavescens* Ait [14,15]. Ma exhibited multipharmacological effects and biological properties and was used to treat inflammation, tumor, virus, fibrosis, arrhythmia, asthma, and growth of human lung cancer cell migration [16–18]. Ma activited Akt/mTOR signaling pathway on anticancer property [19,20], enhanced inhibitive effects of PI3K/Akt on cell proliferation [21], and induced apoptosis and autophagy via inhibiting PI3K/Akt/mTOR pathway [22]. Additionally, Ma exerted an antitumor effect through apoptosis and autophagy by inhibiting the phosphorylation of Akt, mTOR, and their downstream substrates p70S6K and 4EBP1 [16]. Instead, the activation of eIF4E was regulated by the inhibition of 4E-BP1 [16]. Studies on the protective effects of Ma on acute MH induced by isoproterenol (ISO) via Akt/mTOR/p70S6K/eIF4E signaling pathway are lacking.

**Fig. 1.**
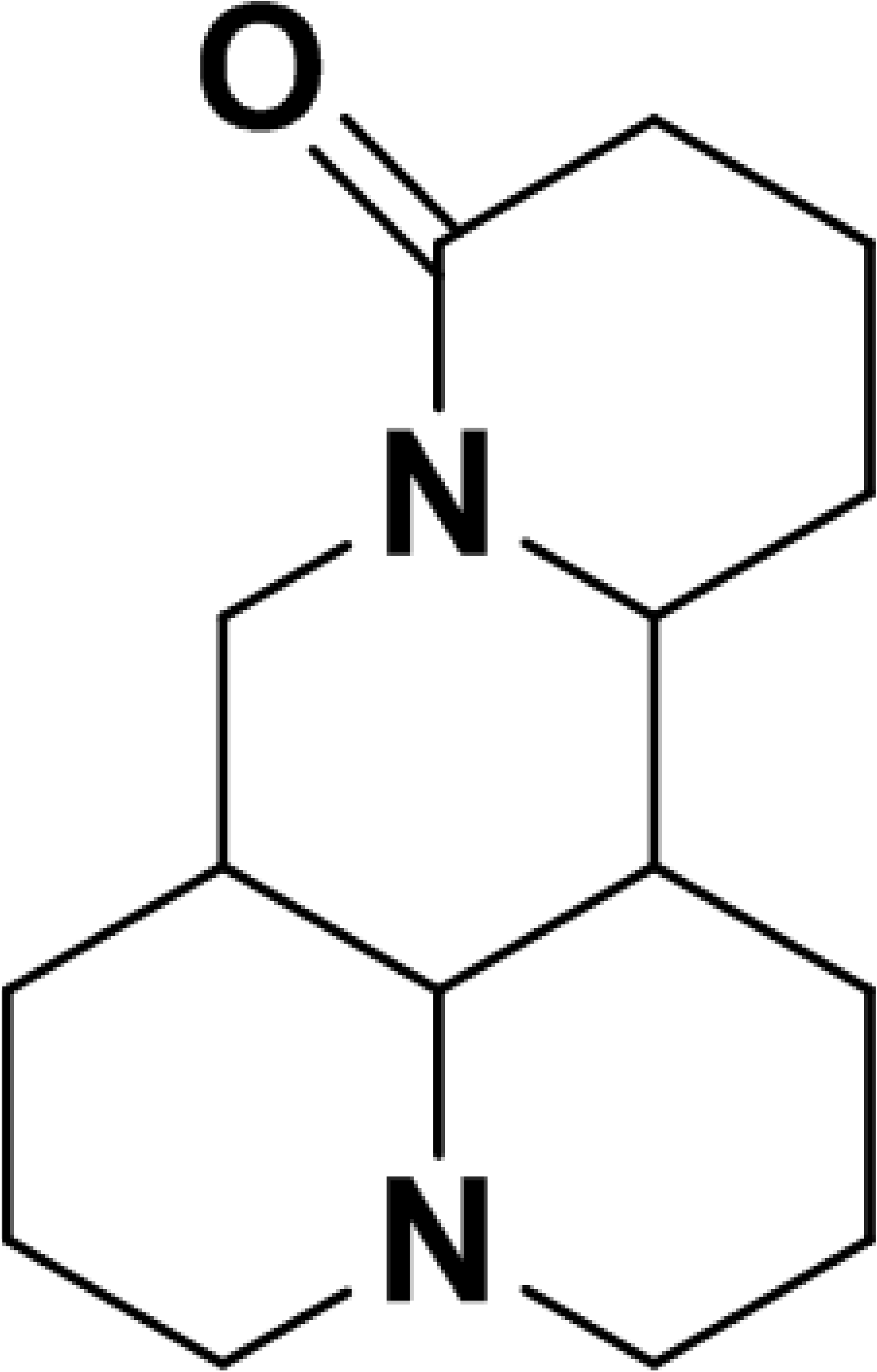
Chemical structure of matrine.

The purpose of this study was to investigate the relationship between Ma on acute MH and Akt/mTOR/p70S6K/eIF4E signaling pathway.

## MATERIALS AND METHODS

### Chemicals and antibodies

Isoproterenol hydrochloride (ISO, 99.9% purity) was purchased from Tokyo Kasei Kogyo Co., Ltd (Tokyo, Japan). Ma (99.8%) was bought from Zijinhua Pharmaceutical Co., Ltd (Ningxia, China), and propranolol was from Sigma Aldrich (≧98.5%, Mainland China). Control saline was purchased from Dajia Pharmaceutical Co., Ltd (0.9%; Guangdong, China). The primary antibodies were as follows: Akt, p-Akt (Ser473), mTOR, p-mTOR (Ser2448), p70S6K and p-p70S6K (Thr389) (Cell Signaling Technology, Inc., Shanghai, China); eIF4E (Abcam, Cambridge, Britain); β-actin (ZSGB-Bio ORIGENE, Beijing, China); goat antimouse secondary antibody and goat antirabbit secondary antibody (Thermo Fisher Scientific, New York, USA). Their primers were synthetized from Invitrogen (Thermo Fisher Scientific, New York, USA).

### Animal and experimental protocol

After obtaining approval from the ethics committee of Ningxia Medical University, the guidelines of the “National Institutes of Health guide for the care and use of Laboratory animals (NIH Publications No. 8023, revised 1978)” were cited in all experiments. Forty-two male Sprague Dawley rats weighing 300±20g were randomly divided into seven groups as follows (n=6): control group (0.9% NaCl); 200 mg/kg Ma group; ISO group (ISO, 85 mg/kg); 10 mg/kg propranolol+ISO group; 200 mg/kg Ma+ISO group; 100 mg/kg Ma+ISO group; and 50 mg/kg Ma+ISO group. Ma and propranolol were intragastricly administrated, and ISO was subcutaneously injected. ISO was carried out for two continuous days once daily. Then, Ma was administered continuously for 7 days.

### Measurements of echocardiography and hemodynamic function

After the final injection of ISO, electrocardiograms [23] were carried out for 24h. The ST segment representing the slow repolarization in the ventricle was dramatically elevated in the ISO-stimulated rats. Then, the symbolic images were captured instantaneously. After 24h of final drug administration, all rats were anesthetized with ethyl carbamate (1.2 g/kg), and then PE 50 cannula was inserted into ventriculus sinister tissues. Parameters [24,25] of the left ventricular systolic pressure (LVSP), left ventricular end-diastolic pressure (LVEDP), left ventricular end-systolic dimension (LVESD), left ventricular end-diastolic dimension (LVEDD), left ventricular anterior wall thickness (LVAWT), and left ventricular posterior wall thickness (LVPWT) were measured. LVSP and LVEDP were measured using the millar conductance catheter. Maximal LVEDD was measured at the time of the QRS deflection, and minimal LVESD was measured at the time of end systole. Fractional shortening (FS) as a measure of systolic function was calculated as FS(%)= [(LVEDD−LVESD)/LVEDD]×100% [25]. Ejection fraction (EF), showing the relationship between drug and the number of contractile segments [26], was calculated as EF(%) = [(LVEDV−LVESV)/LVEDV]×100% [27].

### Myocardial tissue and histomorphology

Ventriculus sinister tissues taken out from all rats were washed by physiological saline and weighed to obtain the cardiac hypertrophy index. Then, apex cordis was sealed up in 10% formalin. Other ventriculus sinister sections were pestled by a grinding apparatus (Schwingmühle TissueLyser 2, Qiagen, Germany) and stored at −80°C for detection. After dehydrating by gradient ethanol, apex cordis was dipped and embedded by wax and then was sectioned for 4 μm. Tissues were placed into a constant temperature incubator at 60±1°C for 20 min after preheating at 48°C. When obtaining rid of wax, section was stained by hematoxylin and eosin (H&E) [28]. H&E tissue images collected using a 40× objective lens were observed by doctors of pathology. The extent of damage in the myocardial tissue was evaluated by the degree of myocardial fibrosis and myocardial hypertrophy, level of interstitial edema, and area of inflammatory cell infiltration.

### Western blot

Total proteins were extracted using protein extraction reagents (Roche Diagnostics Ltd., Shanghai, China) from the myocardial tissue of rats. To determine the total levels and phosphorylation status of Akt, mTOR, p70S6K, and the eIF4E activities, the equivalents of 50 mg of proteins were resolved by 8%–12% sodium dodecyl sulfate polyacrylamide gel electrophoresis and analyzed by protein immunoblotting. Proteins were transferred into polyvinylidene difluoride (PVDF) membranes, blocked with Tris-buffered saline, 0.1% Tween 20 (TBST) containing 5% dried skimmed milk for 2h on a table concentrator [29]. Then, the PVDF membranes were probed with appropriate primary antibodies (Akt, 1:7000; p-Akt, 1:300; mTOR, 1:500; p-mTOR, 1:500; p70S6K, 1:500; p-p70S6K, 1:300; eIF4E, 1:2000, and β-actin, 1:1000) at 4°C overnight [30]. After washing with 5% TBST four times, the membranes were incubated with peroxidase-conjugated secondary antibodies [31] for 2h at room temperature. Signals were visualized by scanning densitometry with the ChemiDic™ MP Imaging System (Bio-Rad Laboratories, Hercules, CA, USA), and the band densities, expressed as relative integrated intensity, were quantified with Quantity-One software (Bio-Rad Laboratories, America). Sample in western blot was triplicate.

### Detection of mRNA by real-time polymerase chain reaction (RT-PCR)

RT-PCR, which was performed with LightCycler 480 SYBR Green І Master Gene Expression Assays (Roche Molecular BiocheMHcals, Germany), is quantitated for the expression of β-actin/Akt/mTOR/p70S6K/eIF4E by employing the 2^−ΔΔCT^ value models [31]. The ΔCT values were expressed as Mean±SD, whereas 2^−ΔΔCT^ was used to analyze the mRNA expression levels. Internal controls were served by the RT-PCR results of β-actin. The primers used in the experiments were designed by oligo (dT) first strand primer. Total RNA was extracted by 1mL Trizol (Roche Molecular Biochemicals, Germany), which was added to a 2 mL polypropylene microcentrifuge tubes containing 0.1g myocardial tissue. Then, the tubes were shaken at 23 rpm for 8 min. Chloroform of 0.3 mL was added to each tube centrifuged at 12,000 rpm at 4°C for 15 min, and the aqueous phase was pipetted into the other clean tubes. After placing 0.5 mL isopropanol into the aqueous phase, the mixture was vortexed at 3000 rpm at 4°C for 1 min and incubated at room temperature for 10 min, and then was centrifuged at 12,000 rpm at 4°C for 10 min. Alcohol of 0.5mL was used for washing the rest of the isopropanol. Finally, we added 30 μL RNase-free and DNase-free water and detected the RNA concentration using 2 μL solution. First strand cDNA was synthesized using PrimeScript™ RT reagent kit with gDNA Eraser (Takara, Dalian, China). After cDNA synthesis, the mRNA expression levels of Akt, p-Akt, mTOR, p-mTOR, p70S6K, and eIF4E were estimated by RT-PCR (LightCycler 480 II, Roche, Switzerland) using the SYBR Green Master.

### Statistical Analysis

All data were represented as Mean±SD, and confidence interval was 95% in our experiment. The One-way ANOVA was used for analyzing the data between the different groups. Statistical results were performed with SPSS (version 22.0, IBM, Armonk, NY, USA) for windows. P<0.05 was considered statistically significant.

## RESULTS

A model of acute MH, induced by ISO rather than Ma, was observed from the results of echocardiography and pictures of pathological section. As shown in Fig.2, myocardial organization structures of the 200 mg/kg Ma and control groups were both clear-cut, which showed no essential effects on myocardial tissues. LVAWT and LVPWT in the ISO group increased, whereas coronary flow decreased. Furthermore, myocardium was incrassated successfully. Moreover, grain of myocardial tissue in the 10 mg/kg propranolol+ISO group was more orderly than that in the ISO group. These findings strongly suggested that MH was induced by ISO.

**Fig. 2.**
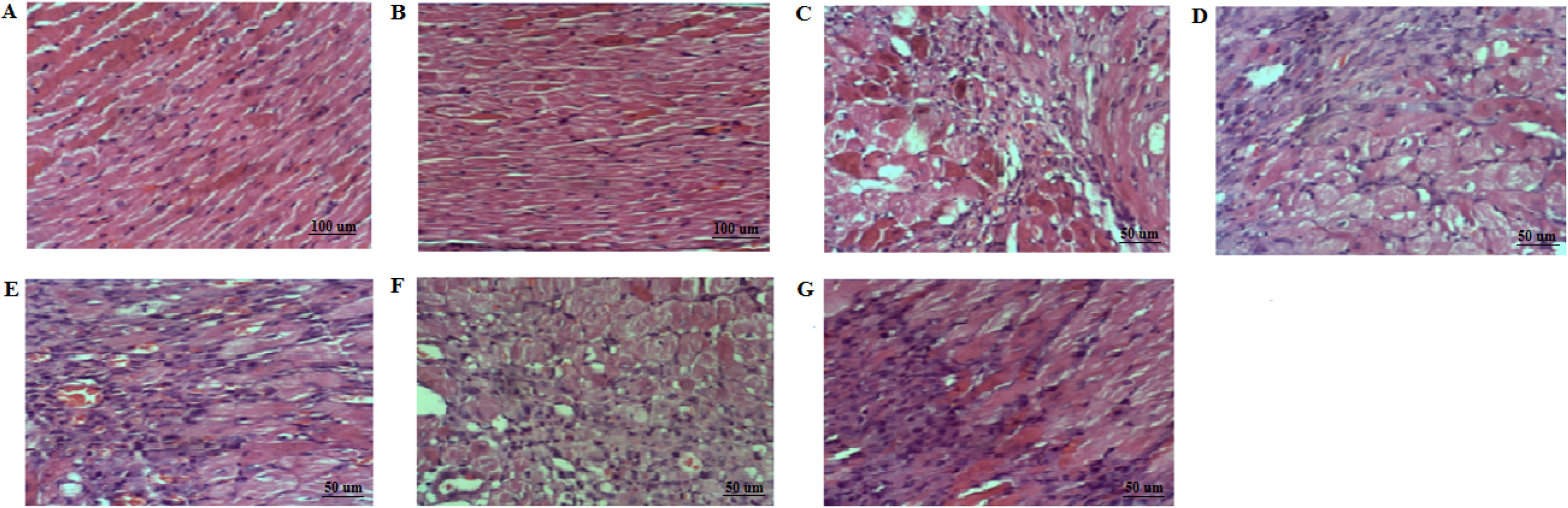
Effects of Ma treatment on histological alterations in the left ventricular myocardial hypertrophy tissue. A: The control group; B: the 200 mg/kg Ma group; C: the ISO (85 mg/kg) group; D: the 10 mg/kg propranolol+ISO group; E: the 50 mg/kg Ma+ISO group; F: the 100 mg/kg Ma+ISO group; and G: the 200 mg/kg Ma+ISO group.

### Symptoms of myocardial hypertrophy alleviated by Ma

The extent of MH alleviated by different doses of Ma is shown in Fig.2. Myocardial infarction size was reduced, and histopathological changes were alleviated in the 200, 100 and 50 mg/kg Ma+ISO groups. Myocardium grains were disordered in the ISO and 50 mg/kg Ma+ISO groups. However, tissue arrangement of the 100 and 200 mg/kg Ma+ISO groups were minimally regular. Histopathological changes of the 200 and 100 mg/kg Ma groups were significantly different from that of the ISO group, whose results were similar to that of the 50 mg/kg Ma+ISO group. Ma exerted curative effects to alleviate the MH symptom.

### Evaluation of echocardiography

Post-pachyntic cardiac dysfunction of MH rats was improved. The results of the control and 200 mg/kg Ma+ISO groups were similar. However, the lacunae vasorum obviously expanded, and the thickness of the vascular wall decreased after treatments with the 200 and 100 mg/kg Ma+ISO than that in the ISO group (Fig.3). The ventricle repolarized, and myocardial systolic function recovered slowly after Ma treatment. These data further confirmed that Ma potently ameliorated the extent of myocardial damage.

**Fig. 3.**
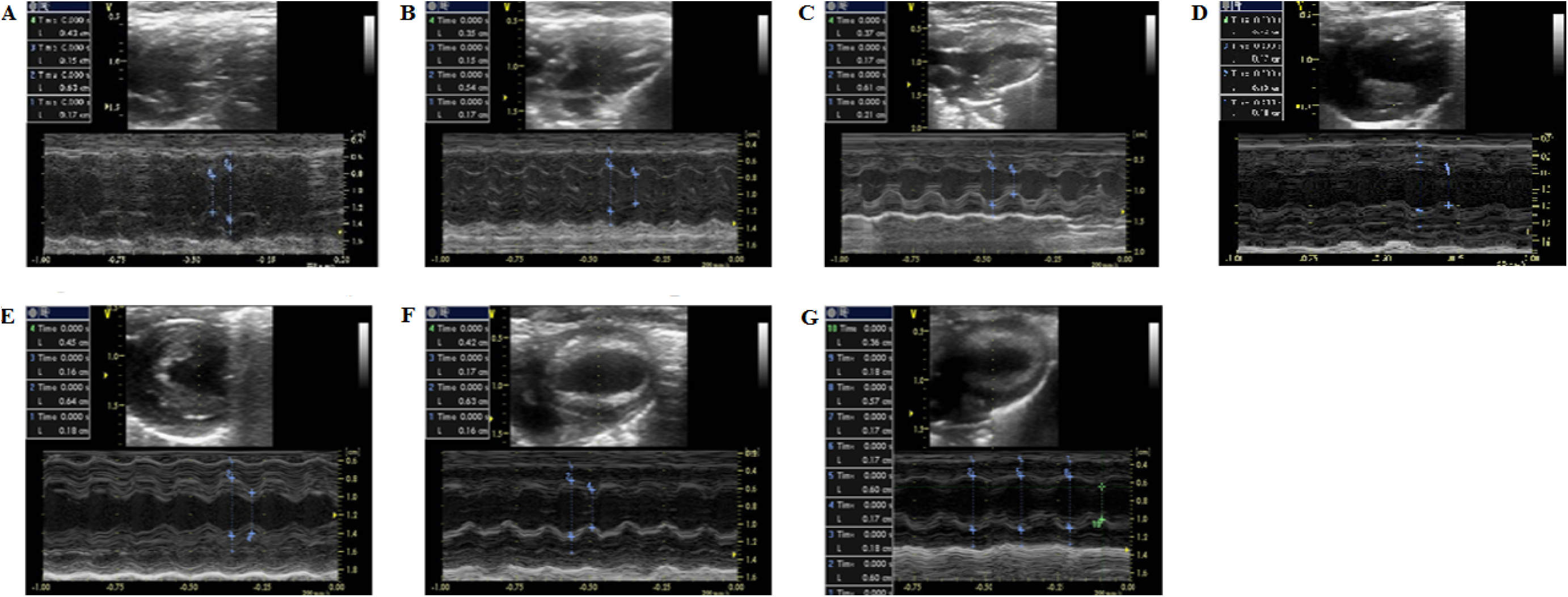
Regions of the left ventricle captured by electrocardiograms in the center position in each segment. A: The control group; B: the 200 mg/kg Ma group; C: the ISO (85 mg/kg) group; D: the 10 mg/kg propranolol+ISO group; E: the 50 mg/kg Ma+ISO group; F: the 100 mg/kg Ma+ISO group; G: and the 200 mg/kg Ma+ISO group.

### Hemodynamic parameters

Cardiac function was assessed using echocardiography. To determine the Ma cardioprotection on MH injury, hemodynamics was evaluated in rats treated with or without Ma. LVAWT and LVPWT of the ISO group were all higher than that of the control and 200 mg/kg Ma groups as shown in Table 1. LVAWT and LVPWT were 1.25- and 1.27-fold higher in the ISO group than that in the control group (P<0.000, 0.000), whereas the results of the 200 mg/kg Ma and control groups were similar (P>0.05). Meanwhile, LVAWT and LVPWT of the 200, 100 and 50 mg/kg Ma+ISO groups significantly decreased compared with that of the ISO group (P<0.05). Above results of the 10 mg/kg propranolol+ISO group were different with that of the ISO group (P=0.047, 0.047). LVESD and LVEDD showed no significance among each group. Meanwhile, for FS and EF, the results of the 100 and 50 mg/kg Ma+ISO groups close to the control group were lower than that of the ISO group. These results revealed that Ma reduced the left ventricular anterior and posterior chamber wall thickness, expanding the left ventricular vessels.

**Table 1.** Hemodynamics parameters of seven treatments in acute myocardial ischemia induced by ISO in rats (Mean ± SD).

|  | LVAWT (mm) | LVPWT (mm) | LVESD (mm) | LVEDD (mm) | FS (%) | EF (%) |
| --- | --- | --- | --- | --- | --- | --- |
| Control group | 0.16 ± 0.01 | 0.15 ± 0.01 | 0.40 ± 0.03 | 0.61 ± 0.04 | 33.00 ± 6.09 | 60.20 ± 8.57 |
| Matrine (200 mg/kg) | 0.15 ± 0.02 | 0.16 ± 0.02 | 0.39 ± 0.04 | 0.61 ± 0.05 | 36.24 ± 4.39 | 64.79 ± 5.95 |
| ISO (85 mg/kg) | 0.20 ± 0.01<sup>$\Delta\Delta$</sup> | 0.19 ± 0.02<sup>$\Delta\Delta$</sup> | 0.41 ± 0.06 | 0.64 ± 0.03 | 35.98 ± 6.94 | 63.90 ± 9.71 |
| Propranolol (10 mg/kg) | 0.19 ± 0.02 | 0.16 ± 0.02 | 0.41 ± 0.04 | 0.64 ± 0.06 | 33.98 ± 3.75 | 61.67 ± 5.02 |
| Matrine (50 mg/kg) + ISO | 0.17 ± 0.00<sup>**</sup> | 0.16 ± 0.01<sup>**</sup> | 0.41 ± 0.03 | 0.62 ± 0.03 | 34.49 ± 2.04 | 62.57 ± 2.94 |
| Matrine (100 mg/kg) + ISO | 0.17 ± 0.02<sup>**</sup> | 0.16 ± 0.01<sup>**</sup> | 0.42 ± 0.06 | 0.63 ± 0.05 | 34.32 ± 7.16 | 61.68 ± 10.37 |
| Matrine (200 mg/kg) + ISO | 0.16 ± 0.02<sup>**</sup> | 0.16 ± 0.02<sup>**</sup> | 0.41 ± 0.04 | 0.65 ± 0.03 | 37.48 ± 4.82 | 66.08 ± 6.44 |

LVAWT = Left ventricular anterior wall thickness; LVPWT = Left ventricular posterior wall thickness; LVESD = Left ventricular end-systolic dimension; LVEDD = Left ventricular end diastolic dimension; FS = Fractional shortening; EF = Ejection fraction; ISO = isoproterenol; vs control: <sup>$\Delta$</sup> $P$ <0.05, <sup>$\Delta\Delta$</sup> $P$ <0.01; vs ISO group: <sup>\*</sup> $P$ <0.05, <sup>\*\*</sup> $P$ <0.01.

### Protein expression and mRNA expression

As shown in Figs.4, protein expression levels of Akt and p-Akt (Ser473) in the control group were 2.10- and 1.89-fold higher than that in the ISO group (P<0.01). The results of Akt and p-Akt in the 200 and 100 mg/kg Ma+ISO groups were 1.57- and 1.81-fold and 1.49 and 1.62-fold higher than that in the ISO group, respectively. Meanwhile, the values were 1.80- and 1.73-fold higher in the 10 mg/kg propranolol+ISO group than that in the ISO group. In terms of mRNA expression of Akt in the 200, 100, and 50 mg/kg Ma+ISO groups, no significant difference was observed compared with that of the ISO group (Fig.3). Moreover, the levels of mRNA expression among the seven groups reached no difference.

**Fig. 4.**
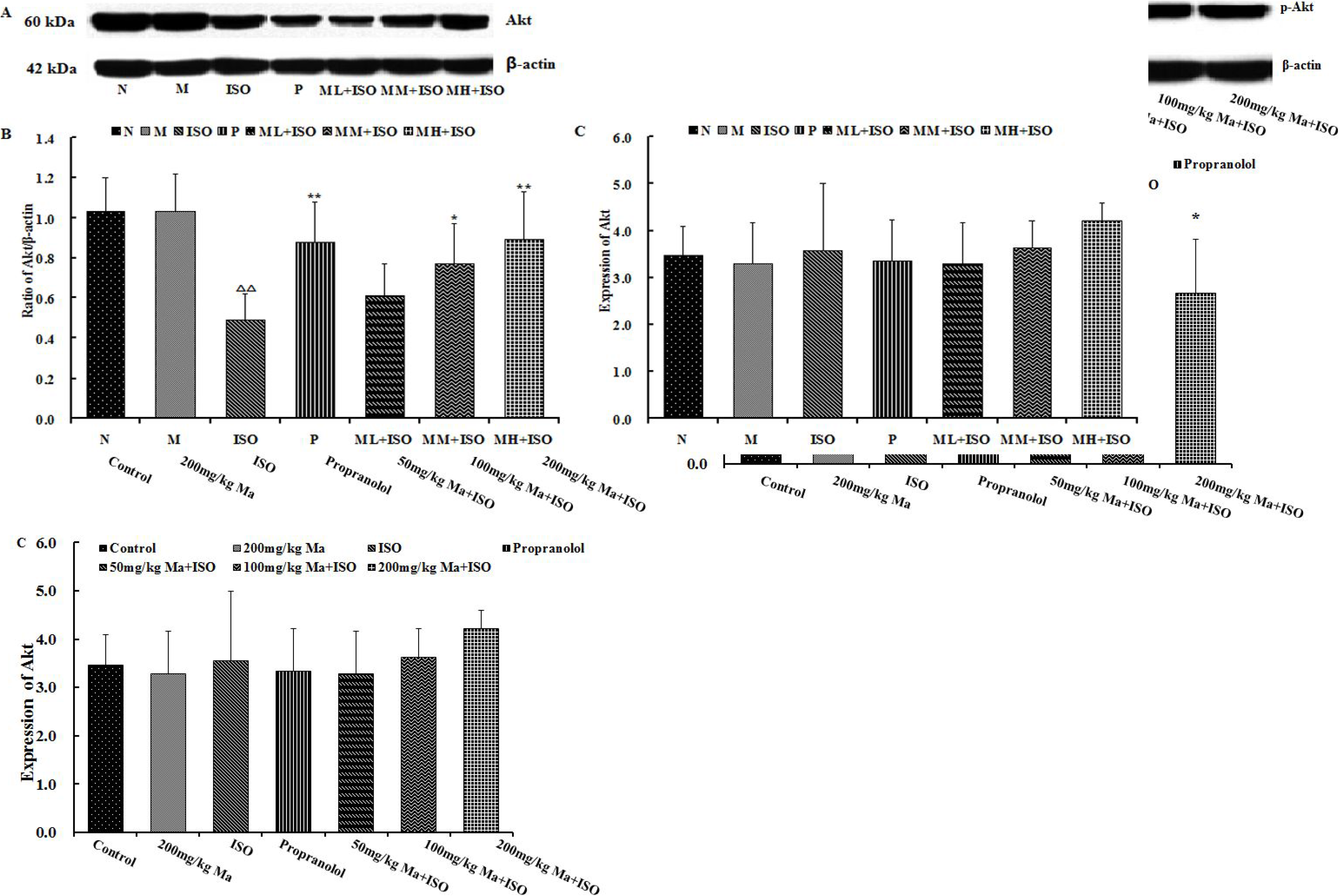
Effects of Ma on the protein expression of Akt and p-Akt represented by Western blot and statistical analysis of the gray value and on the mRNA expression of Akt represented by statistical analysis of the 2^−ΔΔCT^ value in the left ventricular myocardial hypertrophy. A: The control group; B: the 200 mg/kg Ma group; C: the ISO (85 mg/kg) group; D: the 10 mg/kg propranolol+ISO group; E: the 50 mg/kg Ma+ISO group; F: the 100 mg/kg Ma+ISO group; and G: the 200 mg/kg Ma+ISO group. Data are expressed as the mean ± standard deviation (n=6). ^Δ^=vs. the control group, ^Δ^P<0.05, ^ΔΔ^P<0.01; ^*^=vs. ISO group, ^*^P<0.05, ^**^P<0.01.

Protein expression levels of mTOR and p-mTOR (Ser2448) in the ISO group were 2.95- and 7.5-fold higher than that in the control group and continue to increase (Figs.5). Moreover, compared with the ISO group, the expression trends of mTOR and p-mTOR in the 200, 100, and 50 mg/kg Ma+ISO groups were significantly downregulated.

**Fig. 5.**
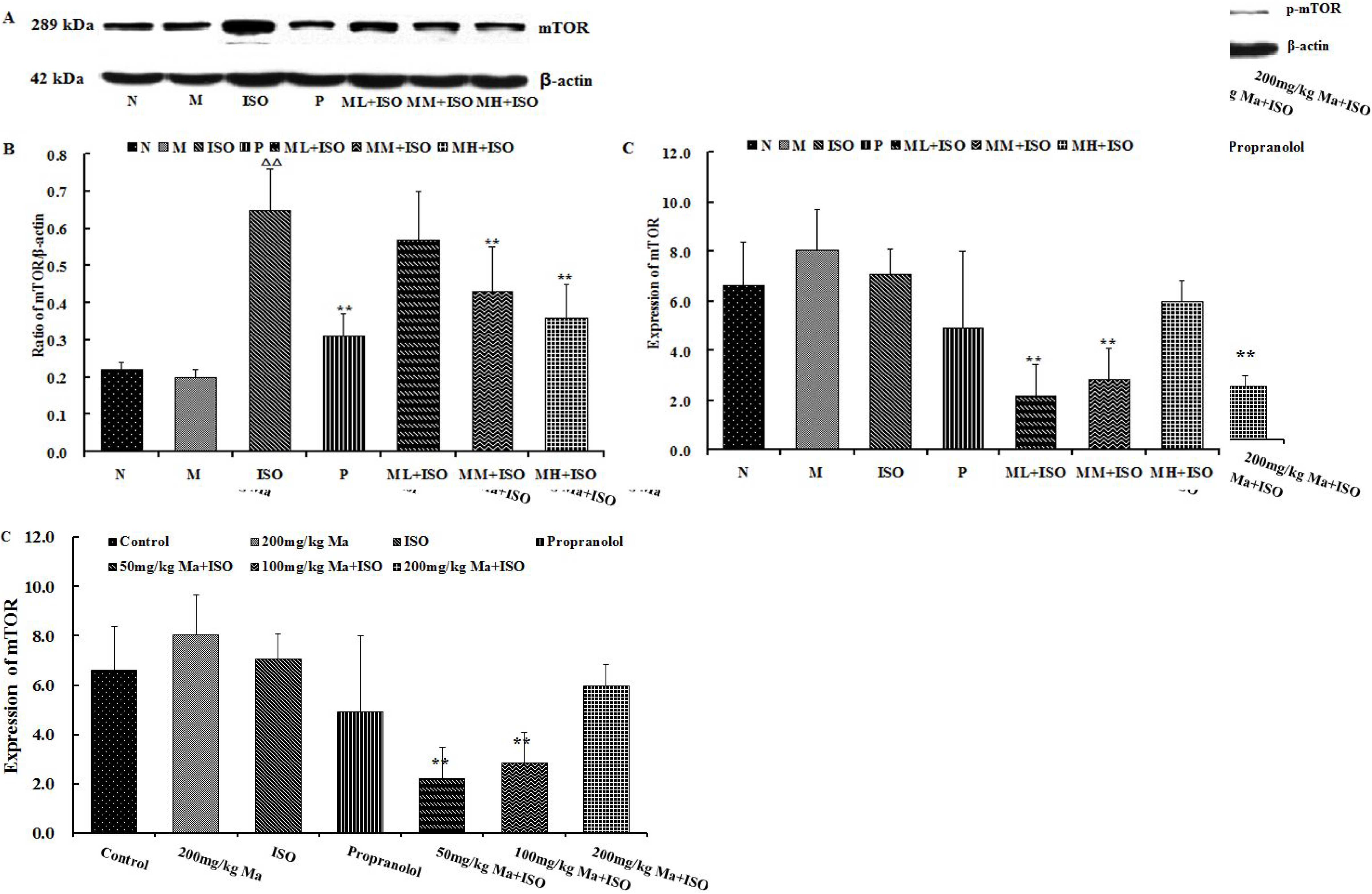
Effects of Ma on the protein expression of mTOR and p-mTOR represented by Western blot and statistical analysis of the gray value and on the mRNA expression of mTOR represented by statistical analysis of the 2^−ΔΔCT^ value in the left ventricular myocardial hypertrophy. A: The control group; B: the 200 mg/kg Ma group; C: the ISO (85 mg/kg) group; D: the 10 mg/kg propranolol+ISO group; E: the 50 mg/kg Ma+ISO group; F: the 100 mg/kg Ma+ISO group; G: the 200 mg/kg Ma+ISO group. Data are expressed as the mean±standard deviation (n=6). ^Δ^=vs. the control group, ^Δ^P<0.05, ^ΔΔ^P<0.01; ^*^=vs. ISO group, ^*^P<0.05, ^**^P<0.01.

However, due to the SD in control group is large, no statistical difference was observed between ISO group and control group on mRNA expression of mTOR (Fig.5). mRNA expression of mTOR in the 100 and 50 mg/kg Ma+ISO groups showed a significant difference compared with that of the ISO group (P=0.003, P=0.001). There is no significant difference in the 200 mg/kg Ma+ISO group and control group on protein and mRNA expression of mTOR.

Protein expression of p70S6K, p-p70S6K (Thr389) and eIF4E was reduced in the 200, 100, and 50 mg/kg Ma+ISO groups according to the ISO group (P<0.05) as shown in Figs.6 and 7. Moreover, mRNA expression of p70S6K in the 100 mg/kg Ma+ISO group was close to that of the control group (Fig.6). mRNA expression of p70S6K and eIF4E in the ISO group were 1.90- and 6.38-fold higher compared with that in the 100 mg/kg Ma+ISO group. The value of the 200 mg/kg Ma+ISO group was close to that of the control group. These findings testified that Ma positively regulated the protein expression of Akt and p-Akt, thereby negatively regulating mTOR/p70S6K/eIF4E signaling pathway on myocardial I/H.

**Fig. 6.**
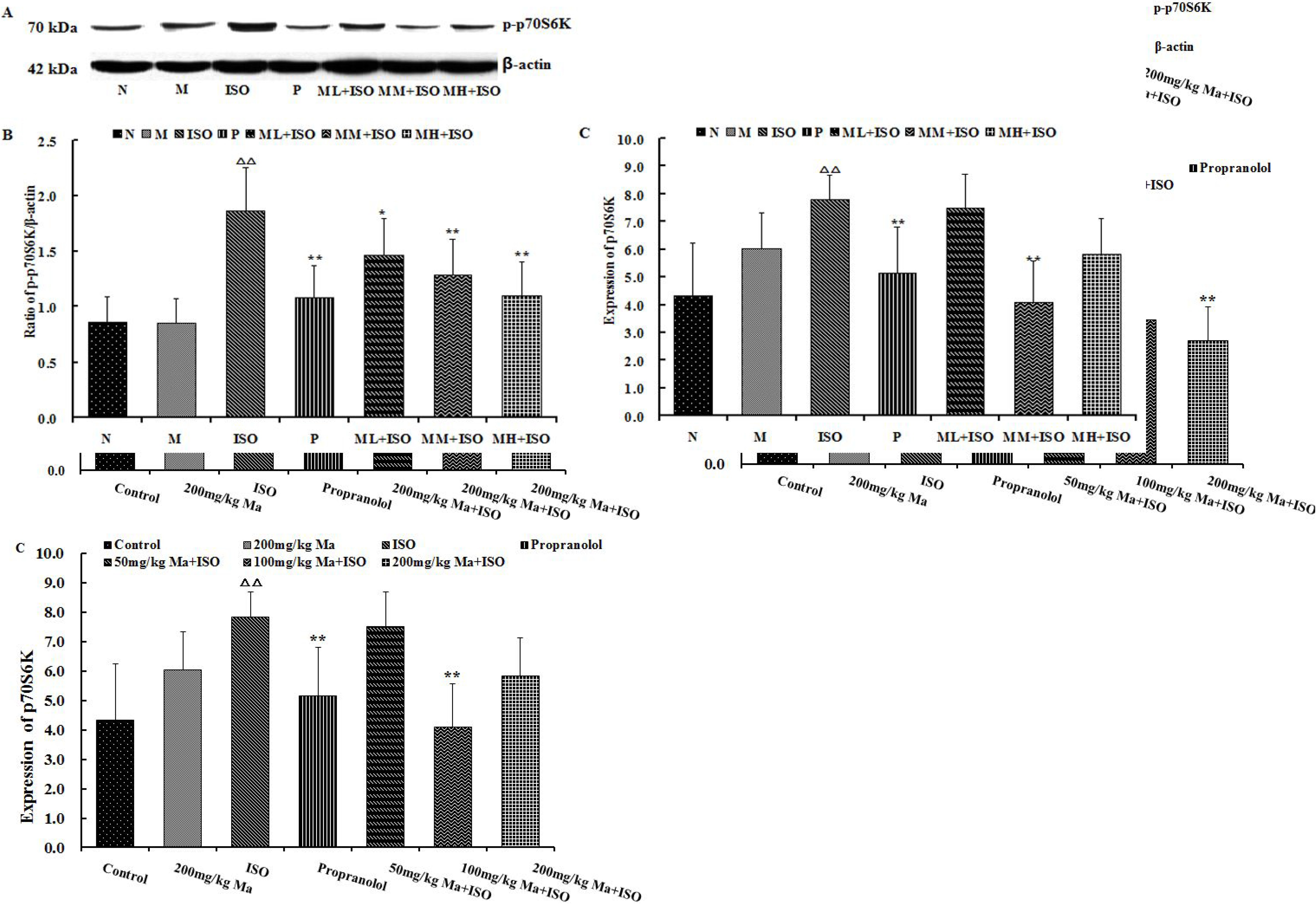
Effects of Ma on the protein expression of p70S6K and p-p70S6K represented by Western blot and statistical analysis of the gray value and on the mRNA expression of p70S6K represented by statistical analysis of the 2^−ΔΔCT^ value in the left ventricular myocardial hypertrophy. A: The control group; B: the 200 mg/kg Ma group; C: the ISO (85 mg/kg) group; D: the 10 mg/kg propranolol+ISO group; E: the 50 mg/kg Ma+ISO group; F: the 100 mg/kg Ma+ISO group; and G: the 200 mg/kg Ma+ISO group. Data are expressed as the mean±standard deviation (n=6). ^Δ^=vs. the control group, ^Δ^P<0.05, ^ΔΔ^P<0.01; ^*^=vs. ISO group, ^*^P<0.05, ^**^P<0.01.

**Fig. 7.**
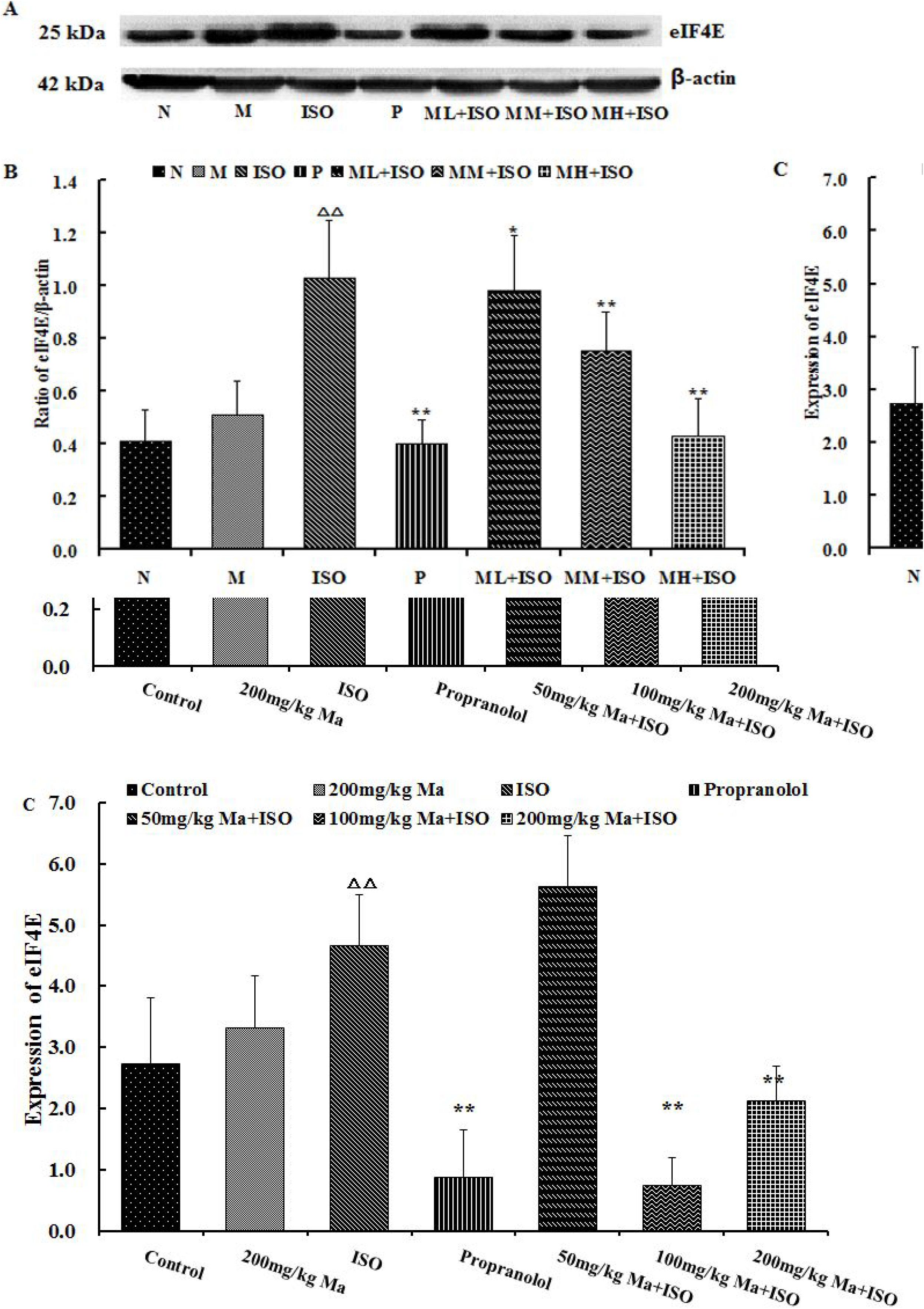
Effects of Ma on the protein expression of eIF4E represented by Western blot and statistical analysis of the gray value and on the mRNA expression of eIF4E represented by statistical analysis of the 2^−ΔΔCT^ value in the left ventricular myocardial hypertrophy. A: The control group; B: the 200 mg/kg Ma group; C: the ISO (85 mg/kg) group; D: the 10 mg/kg propranolol+ISO group; E: the 50 mg/kg Ma+ISO group; F: the 100 mg/kg Ma+ISO group; and G: the 200 mg/kg Ma+ISO group. Data are expressed as the mean±standard deviation (n=6). ^Δ^=vs. the control group, ^Δ^P<0.05, ^ΔΔ^P<0.01; ^*^=vs. ISO group, ^*^P<0.05, ^**^P<0.01.

## DISCUSSION

In the present study, Ma plays an essentially protective effect against MH via Akt/mTOR/p70S6K/eIF4E signaling pathway in rats. Ma could improve the tissue arrangement and alleviate histopathology, showing that Ma exhibits the ability to protect the rat heart against MH. Meanwhile, we investigated the mechanisms responsible for the protective effects of Ma on MH by determining the expression levels of Akt/mTOR/p70S6K/eIF4E factors. Our data indicated that the protective effects of Ma would be associated with regulative expression of Akt/mTOR/p70S6K/eIF4E signaling.

High dose ISO (85 mg/kg) caused MH directly by obviously disordering the myocardium grain [32]. Meanwhile, possible cardiac symptoms or signs would be deterMined by echocardiography in the clinical request. A targeted expansion of echocardiography is likely to increase the detection of clinically important pathology [33]. Echocardiography showed the potential effect of evaluation on tissue viability and the left ventricular contractive function with acute heart failure [34]. The left ventricle structure and dysfunction were reliably performed in MH by echocardiography. Comparable levels of the left ventricular systolic pressure could be explained by the left ventricular wall thickness and contractility sensitivities [35,36]. The weakened ventricular pumping function and myocardial contractility were represented by the increased LVAWT, LVPWT, LVESD, and LVEDD [37,34]. Ventricular wall thickness incrassation induced severe pathological lesion, and ventricular pressure was heightened [38]. FS and EF reflected the myocardial contractility, and when EF was raised, myocardial contractility also increased [37]. The present study, therefore, indicated that histopathological exaMinations were directly used for estimation of LV contractile dysfunction in the MH model. The degree of pachyntic damage was observed by H&E staining, which could be used to identify the histopathological changes associated with MH injury.

Ma exerted neuroprotective effects against pachyntic injury [39,40] and focal cerebral hypertrophy [41], which directly protected neurons and astrocytes. In our experiment, to explore the appropriate dose for treatment, we designed three doses. After treatment with Ma, myocardium grain was ordered, myocardial tissue thickness decreased to different levels, and ventricular pumping function and myocardial contractility improved. In general, Ma exerted a repairing effect on acute MH induced by ISO.

Ma mechanism on acute MH induced by ISO was related to several factors. The research showed that MH was alleviated probably through activating the Akt pathway [42]. mTOR signaling presented both cardioprotective and cardiotoxic effects on ischemic heart disease [43]. Meanwhile, mRNA expression through the Akt/mTOR signaling pathway was regulated [44], and Akt phosphorylation was not altered by hypertrophy [45]. Cardiomyocytes were also protected by activation of PI3K/Akt/mTOR pathway [46]. However, in our experiment, we observed that ISO inhibited protein expression of Akt and p-Akt. Ma upregulated protein expression of Akt and p-Akt but displayed no significant effects on mRNA expression of Akt. ISO also upregulated protein expression of mTOR and p-mTOR and mRNA expression of mTOR, and the tendency was restrained by Ma. It is possible that the inhibition of mTOR signaling pathways were regulated by Akt-independent mechanism. On the other hand, the activation of p70S6K could trigger IRS-1 degradation which then reduces the activation of Akt. Therefore, in the current study, it is also possible that the reduction of p70S6K activated Akt. P70S6K was activated by bone marrow mesenchymal stem cells for improving cardiac contractility and cardiac function [30]. The effects of hypertrophy on the changes in eIF4E assembly and S6K1 phosphorylation were validated [31]. After Ma treatment, the general expression trends of p70S6K and eIF4E were similar to that of mTOR. Akt activity was accelerated by Ma, but activities of mTOR, p70S6K, and eIF4E were inhibited by Ma. This phenomenon was similar to that observed in other studies, thereby showing that both mTOR and p70S6K pathways were inhibited during MH [14, 21]. Ultimately, Ma influenced mRNA expression from p70S6K to its downstream of eIF4E. Ma also showed effects on protein expression from upstream Akt to downstream of eIF4E, whose signaling pathway includes Akt/mTOR/p70S6K/eIF4E. The data showed that Ma would cure MH through the Akt/mTOR/p70S6K/eIF4E signaling.

In general, elevated mRNA levels usually increase protein levels, however, high protein levels usually have low mRNA levels as result of the cell-regulatory mechanism. In terms of cell-balance, cell will be overloaded when the protein level is excessive. In order to maintain the balance, the cell will inevitably reduce the gene transcription because of negative feedback. Meanwhile, eukaryotic gene expression of transcription and translation has a time interval. Gene expression include transcription, post-transcriptional processing, transcriptional product degradation, translation, post-translational processing and modification. The mRNA may have degraded when the protein reaches a peak value, or the protein level is still increasing when the mRNA reaches a peak value. Therefore, the level between protein and gene expression is not consistent.

## CONCLUSION

The present study has demonstrated that Ma plays a protective effect on acute myocardial hypertrophy via Akt/mTOR/p70S6K/eIF4E signaling pathway. Our experiment showed theoretical evidence for myocardial hypertrophy of rat treated by Ma.

## AUTHOR CONTRIBUTION

All authors performed experiments. Hongwan Dang and Wenping Zhang designed the study; Shijie We and Xiaoying Yang contributed new reagents or analytic tools; Hongwan Dang, Wenping Zhang and Hui Ma analyzed data and prepared the manuscript.

## ACKNOWLEDGEMENTS

We acknowledge Institute of Clinical Pharmacology, Department of Pharmacy, General Hospital of Ningxia Medical University (Ningxia, Republic of China).

